# Anti-isoDGR Antibody Inhibits Atherosclerosis Induced by Western Diet in ApoE^-/-^ mice

**DOI:** 10.1101/2025.06.09.658749

**Authors:** Pazhanichamy Kalailingam, SoFong Cam Ngan, Xavier Gallart-Palau, Aida Serra, Arnab Datta, Toh Hean Ch’ng, Evangelia Litsa Tsiani, Panagiota Klentrou, Raj N. Kalaria, Neil E. McCarthy, Dominique de Kleijn, Siu Kwan Sze

## Abstract

**Background:** Degenerative protein modifications (DPMs) accumulate with aging and can alter biomolecule structure and function, including via spontaneous conversion of Asn-Gly-Arg (NGR) to isoAsp-Gly-Arg (isoDGR) motifs that can bind integrins and drive chronic inflammation.

Since isoDGR-modified extracellular matrix proteins are enriched in atherosclerosis and have been associated with rupture-prone plaque characteristics, we hypothesized that antibody neutralisation can inhibit key pathological features including atherosclerotic vascular plaque formation and metabolic dysfunction.

**Methods:** We first examined *Pcmt1*^-/-^ mice which rapidly accumulate isoDGR due to lack of the corresponding repair enzyme to assess the extent of vascular protein damage. We then treated 6-8 week old atherosclerosis-prone (*ApoE*^-/-^) mice which were fed a high-fat Western diet (WD) with weekly dose of 1mg/kg isoDGR-specific monoclonal antibody (isoDGR-mAb) or isotype-matched control (while on diet) for 2 months duration. A regular chow-fed *ApoE*^-/-^ group served as baseline control. Aortic atherosclerotic burden, plaque composition, systemic inflammation, lipid profiles, hepatic steatosis, and metabolic parameters (indirect calorimetry) were assessed.

**Results:** *Pcmt1*^-/-^ mice displayed extensive isoDGR deposition and degeneration of the aortic wall, linking this DPM to vascular structural damage. In the *ApoE*^-/-^ mice, WD induced large aortic root plaques with abundant isoDGR and macrophage infiltration. IsoDGR-mAb treatment decreased plaque size by ∼30% with reduced lipid and collagen content (*p=0*.*001)*. Furthermore, plaques in treated mice contained significantly fewer CD68+ macrophages that also exhibited limited activation. Systemically, isoDGR-mAb modified lipoprotein profiles by decreasing atherogenic VLDL/IDL/LDL cholesterol (*p=0*.*04)* while slightly increasing HDL, accompanied by a reduction in circulating inflammatory proteins. IsoDGR-mAb also protected against hepatic lipid accumulation which was reduced by ∼60% in treated animals (*p<0*.*001*), with indirect calorimetry confirming ∼30% higher oxygen consumption and energy expenditure without change in food intake or physical activity.

**Conclusion:** We identified isoDGR as a key pathological factor involved in the progression of atherosclerosis. Remarkably, isoDGR neutralization diminished plaque inflammation and improved atherosclerotic plaque stability. Our findings support isoDGR neutralization as a promising therapeutic strategy to mitigate both atherosclerosis and aging-associated metabolic dysfunction.

## INTRODUCTION

Aging is the dominant risk factor for cardiovascular diseases including atherosclerosis^1^, which is associated with accrual of degenerative protein modifications (DPMs) in the vascular matrix^2-5^. In particular, asparagine-glycine-arginine (NGR) sequences in long-lived proteins can spontaneously deamidate to form isoaspartate-glycine-arginine (isoDGR) motifs^6, 7^, which act as pro-inflammatory neo-epitopes ^8-10^. In addition, isoDGR can bind integrins including αvβ3 ^6, 7, 11^, thereby triggering leukocyte adhesion and inflammatory signaling. IsoDGR-modified extracellular matrix components (such as fibronectin, laminin, fibrillin and tenascin) have previously been identified in human atherosclerotic plaques ^5, 12^ and associated with rupture-prone plaque characteristics ^2^, and enhance monocyte adhesion via integrin αvβ3 ^3, 6, 7, 11^. It remains, however, unclear whether isoDGR is involved in the progression of atherosclerotic pathology *in vivo*.

Protein-L-isoaspartate (D-aspartate) O-methyltransferase (*PCMT1*) is an enzyme that repairs isoaspartate residues in proteins^13-15^. A recent whole-genome study identified a 619 bp deletion in *PCMT1* associated with an 8.38-fold increased risk of progressive supranuclear palsy ^16^. This finding highlights *PCMT1*’s potential role in degenerative diseases through impaired protein repair mechanisms. Mice deficient in *PCMT1* (*Pcmt1*^-/-^) cannot repair isoAsp damage, leading to accelerated isoAsp deposition, protein damage, and significantly reduced lifespan (typically ∼1-2 months) ^17^. *Pcmt1*^-/-^ tissues exhibit features of inflammation and contain isoDGR deposits co-located with macrophage infiltration ^4, 9, 18^, suggesting that unrepaired isoDGR may drive inflammation. Importantly, even wild-type mice experience a natural decline in *PCMT1* activity with age and likely accumulate isoDGR over time ^9, 18^, which may contribute to age-associated chronic inflammation.

We recently reported that anti-isoDGR monoclonal antibody (isoDGR-mAb) significantly extends the lifespan in *Pcmt1*^-/-^ mice ^9^. Treated animals displayed lower levels of isoDGR in body tissues and reduced systemic inflammation, with substantially longer survival times compared with untreated littermate controls. This outcome provides proof-of-concept that spontaneous protein damage is not irreversible and can be counteracted by engaging immune clearance mechanisms ^9, 19^. Given these insights, we hypothesized that isoDGR is involved in the progression of atherosclerosis, and that neutralizing this damage motif can reduce the disease progression. For this, we first examined *Pcmt1*^-/-^ mice to assess how unchecked isoDGR accumulation impacts vascular structure. Next, we fed Western Diet (WD) to hyperlipidemic apolipoprotein E knockout (*ApoE*^-/-^) mice ^20, 21^ to test whether isoDGR-neutralizing antibody can reduce progression of atherosclerotic plaque burden and rupture-prone plaque characteristics like macrophage influx. This should not only show a causal role in atherosclerotic plaque progression but identify isoDGR as a potential target in reducing atherosclerotic plaque progression.

## MATERIALS AND METHODS

### Mouse models and diets

*Pcmt1*^-/-^ mice (C57BL/6J background) were used to study effects of unrepaired isoDGR accumulation. These mice were generated by heterozygous intercrosses to lack *PCMT1* and thus accumulate isoAsp/isoDGR damage, as described previously^17^. Wild-type littermates served as controls for *Pcmt1*^-/-^ experiments. Apolipoprotein E knockout mice (*ApoE*^-/-^ C57BL/6J, JAX stock number: 002052) were used for the atherosclerosis studies^21^. At 6-8 weeks of age, *ApoE*^-/-^ mice were fed either standard chow diet (CD) or a high-fat/high-cholesterol Western diet (WD; ∼21% fat, 0.2% cholesterol, TD.88137, Teklad) to induce hyperlipidemia and atherogenesis. Mice remained on their assigned diets for 16 weeks in total. After the first 8 weeks on WD (to establish baseline lesions), WD-fed mice were randomly assigned to receive treatment with isoDGR-mAb or an isotype-matched control (IgG) antibody for the next 8 weeks (while continuing on WD). A separate group of *ApoE*^-/-^ mice maintained on regular CD served as a normolipidemic reference. All mice were housed under 12-hour light/dark cycles with free access to food and water. Animal procedures were conducted in a humane manner and were approved by the NTU Institutional Animal Care and Use Committee (IACUC protocol # ARF-SBS/NIE/LKC-A18016, A19029, A18059).

### Anti-isoDGR antibody treatment

The isoDGR-mAb was developed to specifically recognize the isoDGR motif as previously described by Park *et al*. ^4^. Briefly, mice were immunized with an isoDGR-containing peptide conjugated to a carrier, and hybridomas were screened for antibodies that displayed high-affinity selective binding to isoDGR (but not isoD, NGR, DGR, or RGD sequences). For the treatment group, purified isoDGR-mAb was administered intraperitoneally at 1 mg/kg body weight once weekly. Control mice received an equivalent dose of isotype-matched IgG on the same schedule. Treatments were given during weeks 9-16 of WD. Normal CD-fed control mice did not receive any antibody.

### Tissue collection and histology

At week 16, mice were fasted 4 hours and then deeply anesthetized with isoflurane and euthanized by cardiac puncture (exsanguination). Blood was collected into EDTA tubes for plasma isolation. The arterial tree was perfused with cold phosphate-buffered saline (PBS) through the left ventricle to flush blood, and then dissected for analysis. The heart, liver, and ascending aorta were embedded in OCT compound. Aortic root (including the aortic valve area) was collected as cross-sections (8-10 µm thick) for histological and immunofluorescence analysis. Tissue sections were stained with hematoxylin and eosin (H&E) or Masson’s Trichrome to assess plaque morphology and area, with Oil Red O (ORO) to detect neutral lipid in plaques. *En face* staining of the aortic arch and thoracic aorta with ORO was performed to visualize lipid-rich lesions of the luminal surface. Digital images of stained sections and *en face* aortae were captured for quantitative analysis. Lesion area and ORO-positive area were quantified using ImageJ.

### Immunofluorescence staining

Cryosections of aortic root and liver were fixed in 4% paraformaldehyde, permeabilized (0.1% Triton X-100), and blocked with normal serum. Sections were incubated overnight at 4°C with primary antibodies against isoDGR (using isoDGR-mAb as the probe), macrophage markers (rat anti-F4/80 and rat anti-Mac-2 for plaque macrophages; rabbit anti-CD68 for liver Kupffer cells). After washing, appropriate fluorophore-conjugated secondary antibodies were applied. Nuclei were counterstained with DAPI. Stained sections were mounted and imaged by confocal microscope (Zeiss LSM710) or fluorescence microscope (Nikon Eclipse Ti2). Correlation of isoDGR with cell markers was assessed, and fluorescent signal areas/intensities were quantified using ImageJ software.

### Plasma and liver Analyses

Cytokine concentrations in blood plasma were quantified by LEGENDplex™ multiplex bead assay (Biolegend, San Diego, CA) Using the mouse inflammation panel (IL23, IL1α, IL1β, IL6, IL10, IL12p70, IL17A, IL23, IL27, MCP1, IFNβ, IFNγ, TNFα, and GMCSF). Plasma C-reactive protein (CRP) levels were measured using a mouse-specific ELISA kit (e.g., R&D Systems, MCRP00), according to the manufacturer’s protocol. Plasma lipoprotein profiles were assessed by Fast Protein Liquid Chromatography (FPLC) fractionation of pooled plasma; Total cholesterol in each fraction was quantified using an enzymatic colorimetric assay (e.g., Wako Cholesterol E). VLDL, LDL, and HDL fractions were assigned based on elution profiles and quantified by summing the cholesterol content across corresponding fractions. Liver weight was recorded to gauge hepatomegaly. Sections of liver were stained with ORO to visualize neutral lipid accumulation; the fractional area of ORO-positive staining was quantified as a measure of hepatic steatosis.

### Metabolic cage studies

In the final week of the experiment, mice were placed in metabolic cages for indirect calorimetry (PhenoMaster system) and activity monitoring. Oxygen consumption (VO_2_) and carbon dioxide production (VCO_2_) were recorded over a 48-hour period (with 12:12 h light-dark cycle) along with food and water intake and locomotor activity (beam breaks). Energy expenditure was calculated from gas exchange measurements, and respiratory exchange ratio (RER = VCO_2_/VO_2_) was determined to assess substrate utilization. Body weights were tracked throughout the study.

### Statistical analysis

Data are presented as mean ± standard error of the mean (SEM). For comparisons of three groups (e.g., CD, WD+isotype, WD+isoDGR-mAb), one-way analysis of variance (ANOVA) was used with post-hoc tests (e.g., Tukey’s multiple comparisons) to explore individual group differences. For repeated measures (e.g., metabolic time-course data), two-way ANOVA was applied. For direct comparisons between two groups, two-tailed unpaired *t*-tests were used. Significance was defined as *p* < 0.05.

## RESULTS

### Failure to repair isoDGR protein damage is associated with vascular degradation *in vivo*

Thoracic aortae from six-week-old mice lacking the isoaspartate repair enzyme (*Pcmt1*-/-) revealed structural abnormalities compared to wild-type (WT) controls (Figure 1). WT aortae exhibited the typical wavy pattern of elastic lamellae in the tunica media, whereas *Pcmt1*-/-aortae showed focal thinning and straighter elastic fibers, suggesting compromised vascular elasticity (Figure 1A). Immunofluorescence confirmed extensive accumulation of isoDGR-modified proteins throughout the *Pcmt1*^-/-^ aortic wall, co-localizing with the areas of damage. *Pcmt1*^-/-^ aortic sections showed strong isoDGR immunofluorescence signal in all layers of the vessel (intima, media, and adventitia), whereas age-matched WT aortae displayed minimal isoDGR signal (Figure 1B). The intimal surface of *Pcmt1*^-/-^ aortae appeared undulated and thickened in regions rich in isoDGR, consistent with underlying structural deformation and matrix accumulation. Quantification of fluorescence confirmed significantly greater isoDGR content in *Pcmt1*^-/-^ aortae compared with WT (Figure 1C). Masson’s Trichrome staining of the aorta (Figure 1D) revealed a localized bulging deformity in the vessel wall, resembling an aneurysm, observed in some *Pcmt1*^-/-^ mice. This finding suggests arterial wall weakening associated with loss of *Pcmt1*. Collectively, these findings indicate that failure to repair isoAsp leads to excessive isoDGR deposition accompanied by loss of arterial integrity and severe structural damage. In contrast, WT mice with intact *PCMT1* function maintain low isoDGR levels and normal arterial morphology at the same young age, underscoring the protective role of protein repair in vascular health.

**Figure 1.**
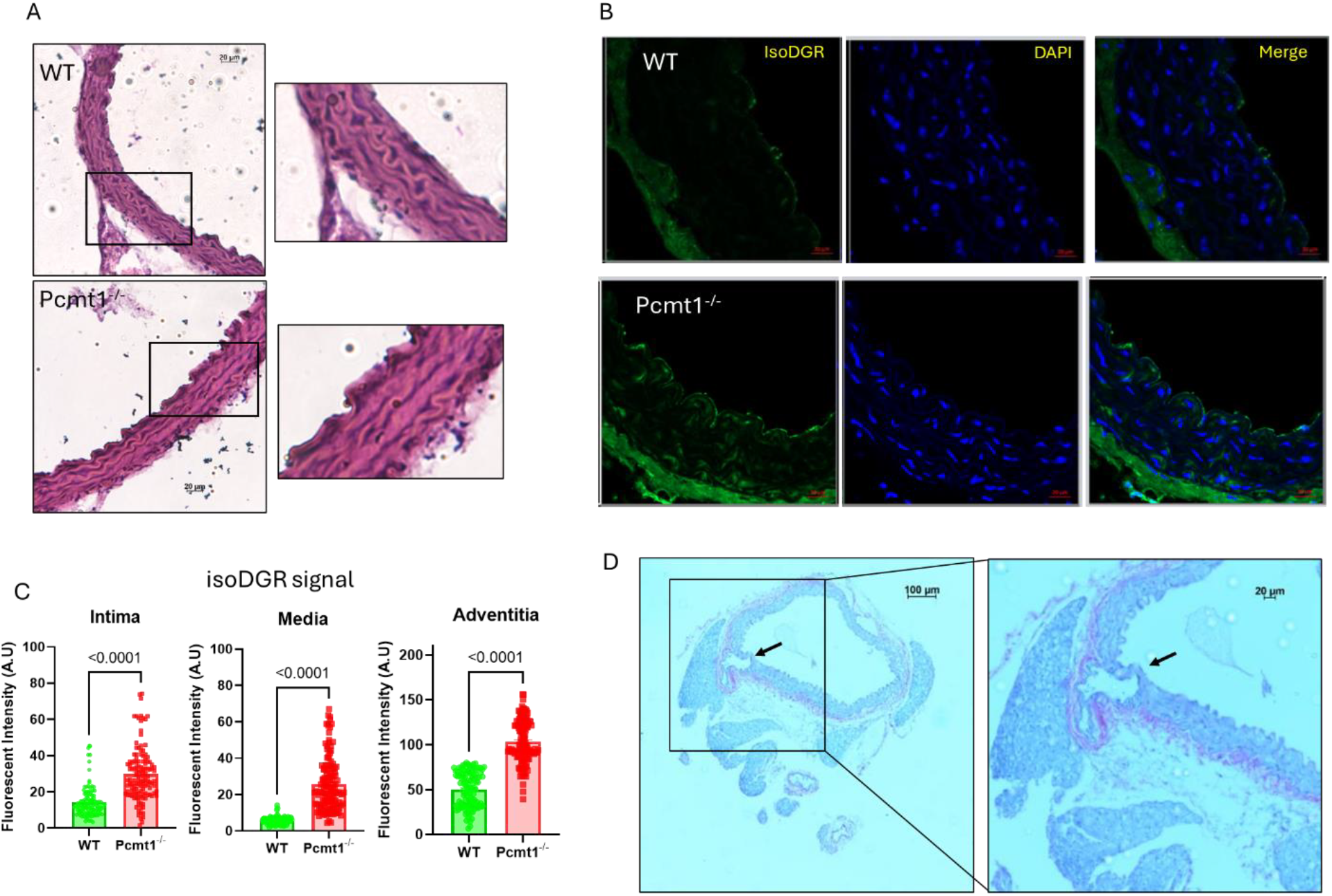
PCMT1 deficiency leads to isoDGR accumulation and aortic wall damage. (A) H&E-stained transverse aortic sections from a wild-type (WT) mouse and a Pcmt1^-/-^ mouse. The Pcmt1^-/-^ aorta exhibits disruption of elastic lamellae and structural deterioration unlike WT tissue. (B) Immunofluorescence staining for isoDGR (green) in aortic sections (nuclei counterstained blue). WT aorta shows minimal isoDGR, whereas Pcmt1^-/-^ aorta exhibits widespread isoDGR deposition. (C) Quantification of isoDGR fluorescence intensity in the aortic wall confirms significantly greater isoDGR accumulation in intima, media, and adventitia from Pcmt1^-/-^ mice. (D) Masson’s Trichrome stain of a Pcmt1^-/-^ aorta highlighting a focal wall outpouching (arrow) with loss of elastic fibers. These findings illustrate that lack of PCMT1-mediated repair causes isoDGR deposition and focal vascular damage. Data are representative of experiments with 3 mice per group (n = 3).

### Anti-isoDGR immunotherapy inhibits WD-induced atherosclerosis

Having established that isoDGR accumulation correlates with vascular degeneration in *Pcmt1*^-/-^ mice, we next examined whether neutralizing isoDGR affects progression of atherosclerosis in the hyperlipidemic *ApoE*^-/-^ model. *ApoE*^*-/-*^ mice on a 16-week WD developed substantial atherosclerotic plaques in the aortic root and arch. WD-fed *ApoE*^*-/-*^ mice treated with isoDGR-mAb displayed markedly lower plaque area than isotype-mAb-treated controls. Aortic root cross-sections (Figure 2A-C) from control isotype-mAb-treated WD-fed *ApoE*^-/-^ mice revealed large plaques with thick intimal lesions, heavy lipid deposition (Oil Red O staining), and collagen-rich areas, whereas isoDGR-mAb-treated mice had significantly smaller plaques with reduced lipid and collagen content (Figure 2A-C). H&E sections showed large, lumen-occluding plaques with necrotic cores in isotype-mAb-treated controls, whereas isoDGR-mAb-treated animals displayed markedly smaller intimal lesions (Figure 2A). Morphometric analysis indicated ∼30% reduction in mean plaque cross-sectional area relative to control WD mice (p < 0.0001), while ORO staining of serial sections highlighted parallel changes in neutral-lipid deposition (Figure 2B). Control plaques contained extensive lipid-laden foam-cell zones, whereas isoDGR-mAb-treated lesions harboured compact lipid cores occupying 30-40% less plaque area (p = 0.003). Masson’s trichrome staining of plaques revealed abundant collagen (blue) in the control lesions, indicating an advanced fibrous composition, while the treated plaque contained markedly less collagen. Histological analysis confirmed that plaques from the control group contained dense, disorganized collagen interwoven with fragmented elastic fibers, which are hallmarks features of advanced, rupture-prone lesions. In contrast, plaques from the treated group displayed a thinner but uniform collagen cap overlaying a markedly smaller lipid core with largely intact elastic laminae. This shift toward organized, surface-focused collagen deposition, with reduced deep-plaque fibrosis, indicates that isoDGR-mAb treatment limits overall plaque area while simultaneously reinforcing cap stability, thereby lowering rupture-prone plaque characteristics without excessive luminal scarring. *En face* staining of the aortic arch for lipid-rich lesions indicated extensive fatty streaks and plaque coverage in isotype-control-treated WD mice, compared with notably fewer and smaller lesions in isoDGR-mAb-treated mice (Figure 2D). Quantitative analysis confirmed that isoDGR neutralization reduced plaque cross-sectional area and *en face* lesion area by ∼30-40% (*p* < 0.0001). This indicates effective inhibition of atherosclerotic progression throughout the vasculature, yielding smaller plaques with less lipid accumulation.

**Figure 2.**
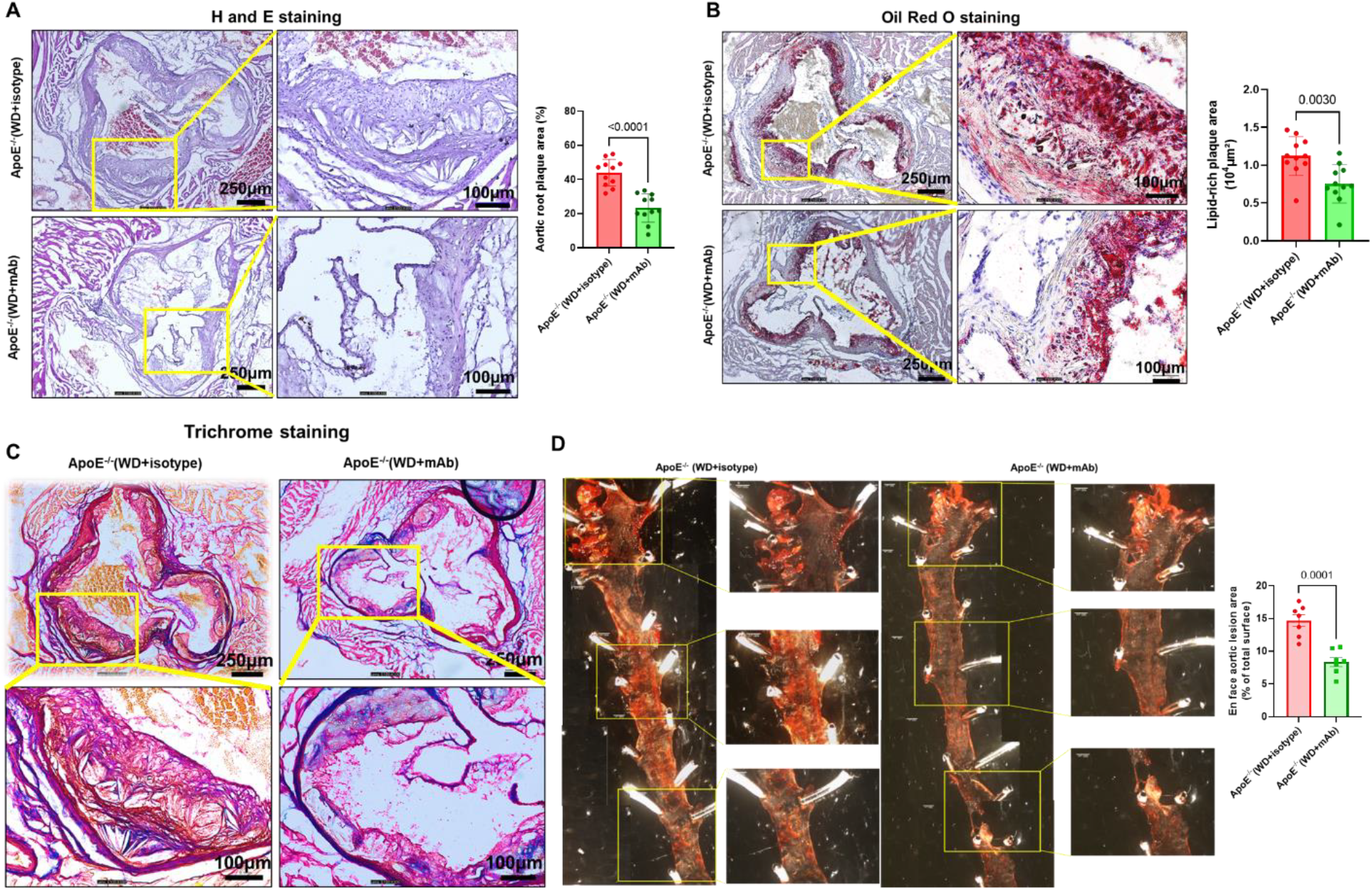
Anti-isoDGR antibody treatment reduces atherosclerotic plaque burden. (A) H&E-stained cross-sections of the aortic root from a WD-fed ApoE^-/-^ mouse treated with isotype control only (upper) or isoDGR-mAb (lower). The control mouse exhibits a large plaque protruding into the lumen, whereas the treated mouse displays a much smaller lesion. (p < 0.0001) (B) Serial sections stained with Oil Red O for neutral lipids highlighted extensive lipid-rich areas (red) in the control plaque, which were markedly reduced in treated plaque. (p = 0.003) (C) Masson’s trichrome staining revealed abundant collagen (blue) in the control lesion, indicating an advanced fibrous plaque, while the treated plaque contained markedly less collagen. Histological analysis shows that plaques from the control group contain dense, disorganized collagen interwoven with fragmented elastic fibers, whereas plaques from the treated group display a thinner and uniform collagen cap overlaying a markedly smaller lipid core with largely intact elastic laminae. (D) En face Oil Red O staining of the aortic arch and thoracic aorta. The control-isotype-treated ApoE^-/-^ aorta displays widespread lipid deposition (red), whereas the isoDGR-mAb treated aorta featured substantially fewer lipid streaks. Overall, isoDGR-mAb therapy significantly reduced plaque size and lipid content in the aorta (p < 0.0001). Data are representative of experiments with 6-12 mice per group (n = 6-12).

### IsoDGR-mAb decreases plaque inflammation and improves structural stability

In addition to reducing plaque size, isoDGR immunotherapy also had profound effects on plaque composition, immune infiltrate, and indicators of structural stability. WD-fed *ApoE*^-/-^ mice that were untreated or received control antibody exhibited highly inflamed plaques, characterized by abundant macrophage infiltration that co-localized with isoDGR deposition. In contrast, plaques from isoDGR-mAb-treated mice displayed markedly less inflammation and more intact normal appearing structural features. Immunofluorescence imaging of aortic root plaques revealed strong isoDGR staining and numerous CD68+ macrophages in control mice (Figure 3A), whereas isoDGR-mAb-treated mice exhibited only faint isoDGR signal and far fewer CD68+ cells in plaques. This qualitative observation was reinforced by co-staining of additional macrophage markers F4/80 and Mac-2, which confirmed that control plaques were densely populated with foam cells, unlike tissues from isoDGR-mAb-treated mice (Figure 3B). Overall, Mac-2 signal in control plaques was particularly intense in areas rich in extracellular lipids (e.g. near the necrotic core), whereas macrophage frequency was far lower in treated plaques, and Mac-2 staining was significantly reduced. Quantitative analysis revealed a marked reduction of isoDGR-modified proteins in atherosclerotic plaques, over 8-fold, following isoDGR-mAb treatment. This was accompanied by a decrease in macrophage infiltration. Specifically, the area of Mac-2+ staining was reduced by more than 50% (p < 0.0001), and the proportion of plaque area occupied by CD68+ cells decreased by over fourfold. These observations demonstrate that neutralizing isoDGR curtailed both the recruitment and activation of macrophages in atherosclerotic lesions. The observed reductions in plaque macrophages are consistent with the concept that isoDGR-modified proteins can act as leukocyte chemoattractants and/or adhesins ^3, 6-11^, thus promoting monocyte recruitment into plaque tissue and local differentiation into CD68+ Mac-2+ foam cells. These results indicate that isoDGR neutralization also decreases inflammatory cell recruitment to vascular lesions.

**Figure 3.**
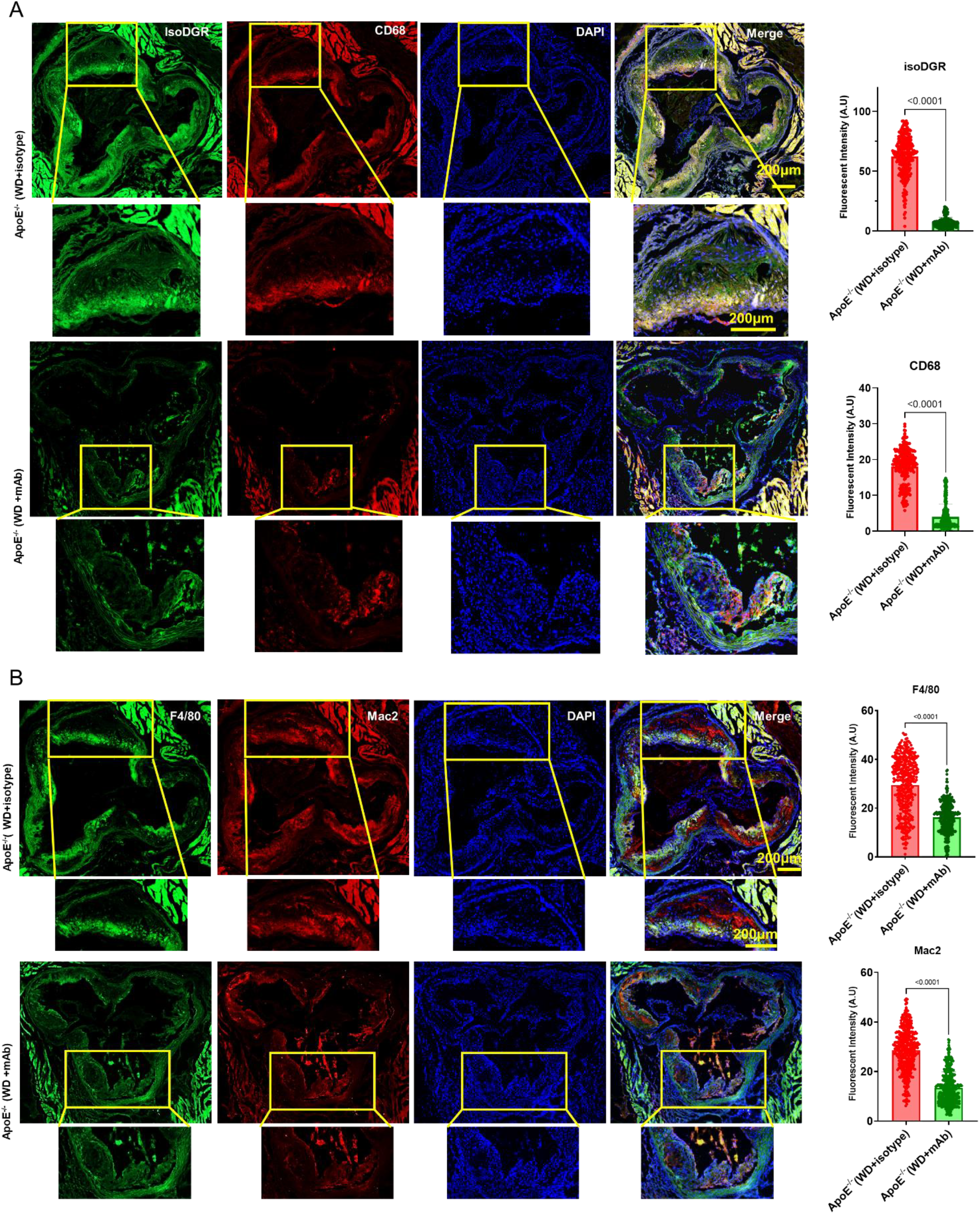
IsoDGR neutralization diminishes plaque inflammation and improves plaque stability. (A) Aortic root lesion immunofluorescence staining for isoDGR (green) and macrophage marker CD68 (red) in WD-fed ApoE^-/-^ mice. Isotype control-treated plaque (left) shows strong isoDGR and CD68 signals which were co-localized (yellow), whereas isoDGR-mAb-treated plaque (right) displayed minimal isoDGR and far fewer CD68+ cells. (B) Dual staining for macrophage markers F4/80 (green) and Mac-2 (red) in plaques. The control plaque is densely infiltrated with F4/80+ cells, many of which were also Mac-2+ (appearing yellow), indicating numerous activated foam cells. In contrast, treated plaque contained markedly fewer macrophages and negligible Mac-2 signal. Results are representative of experiments conducted with 4-6 mice per group (n = 4-6).

### Anti-isoDGR therapy inhibits systemic inflammation and partially restores lipid profile

In addition to isoDGR-induced vascular inflammation, excess lipid intake causes hyperlipidemia, which is known to activate innate immune pathways, elevate pro-inflammatory cytokine expression, and disrupt metabolic homeostasis. We therefore investigated whether the therapeutic effects of isoDGR-mAb extend beyond the vessel wall, potentially impacting systemic inflammation and circulating lipid profiles. We measured circulating cytokines and CRP level in *ApoE*^-/-^ mice fed CD or WD that were treated or not with isoDGR-mAb (WD+mAb) or isotype control. WD feeding increased a broad panel of inflammatory cytokines in blood plasma, including IL-1α, IL-1β, IL-6, TNFα, MCP-1, GM-CSF, IL-12p70, IL-17A, and IL-23, in addition to elevating CRP and decreasing anti-inflammatory IL-10 (Figure 4A). Treatment with isoDGR-mAb reduced most of these cytokines toward CD levels, with significant decreases observed in IL-27, MCP-1, and TNFα as well as CRP. We next examined whether lipid metabolism was similarly impacted by isoDGR-mAb treatment using FPLC fractionation of lipoprotein particles coupled with cholesterol assay (Figure 4B). The WD groups were severely hypercholesterolemic (as is typical for *ApoE*^-/-^ animals on WD), but there were notable improvements in the lipid profile of isoDGR-mAb-treated mice. While WD caused severe hypercholesterolemia, with predominant elevation in atherogenic VLDL/IDL/LDL cholesterol and reduction in HDL, mice receiving isoDGR-mAb showed a significant reduction in total cholesterol and partial rebalancing of VLDL/IDL/LDL content (Figure 4C).

**Figure 4.**
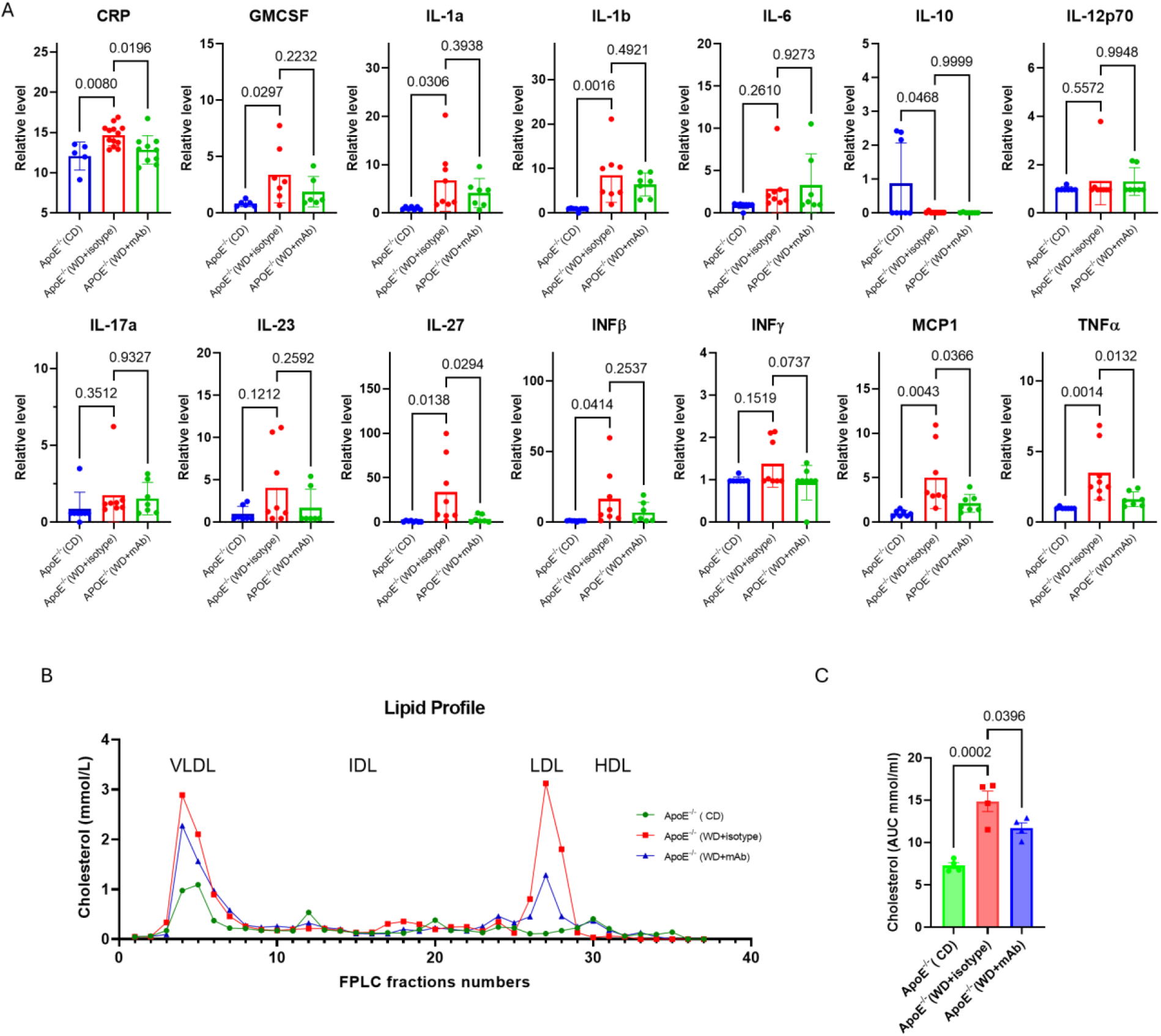
Anti-isoDGR mAb inhibits systemic inflammation and partially restores lipid profile. (A) Plasma cytokines and C-reactive protein (CRP) levels in ApoE^-/-^ mice: WD feeding significantly increased CRP and proinflammatory cytokine levels compared to chow diet, and treatment with isoDGR-mAb significantly reduced CRP, IL-27, MCP-1 and TNFα levels. (B) Plasma cholesterol distribution across lipoprotein fractions (VLDL, LDL, HDL) as measured by FPLC. WD-fed mice (red line) displayed high VLDL/LDL cholesterol peaks relative to chow diet (blue line). IsoDGR-mAb-treated mice (blue line) exhibited lower VLDL/LDL peaks and slight increase in HDL compared to untreated WD, indicating partial correction of lipid profile. (C) Cholesterol levels in plasma FPLC fractions were quantified by calculating area under the curve (AUC) across all fractions. These results suggest that isoDGR-mAb reduces systemic inflammation and influences lipoprotein composition. Representative data shown from experiments involving 4-8 mice per group (n = 4-8).

### IsoDGR-mAb ameliorates WD-induced hepatic steatosis and inflammation

High-fat WD not only caused atherosclerosis in *ApoE*^-/-^ mice but also led to features of metabolic syndrome such as hepatic steatosis (fatty liver) and inflammation (steatohepatitis). Therefore, we further tested if isoDGR neutralization protected against the development of fatty liver and tissue inflammation in WD-fed *ApoE*^-/-^ mice. After 16 weeks of WD, control *ApoE*^-/-^ mice displayed classic signs of diet-induced non-alcoholic fatty liver disease (NAFLD) (Figure 5A). The livers of the mice were enlarged, pale, and greasy in appearance (hepatomegaly, liver ∼6-10% of total body weight vs ∼4-5% for chow diet, Figure 5B) with extensive neutral lipid accumulation in hepatocytes (diffuse Oil Red O staining) (Figure 5C). In striking contrast, isoDGR-mAb-treated mice had much smaller livers (∼20% lower liver weight relative to untreated WD, approaching CD-fed mice) and ∼60% lower hepatic lipid deposition (ORO-positive area, Figure 5D). Histologically, treated livers showed only scattered small lipid droplets, whereas untreated WD livers were filled with large coalescent fat droplets. This finding indicates that blocking isoDGR can protect against diet-induced steatosis. Treated WD livers were still heavier than regular CD diet, indicating that some fat accumulation occurred, but the degree of hepatomegaly was greatly attenuated by antibody treatment. These results indicate partial but significant protection against diet-induced fatty liver.

**Figure 5.**
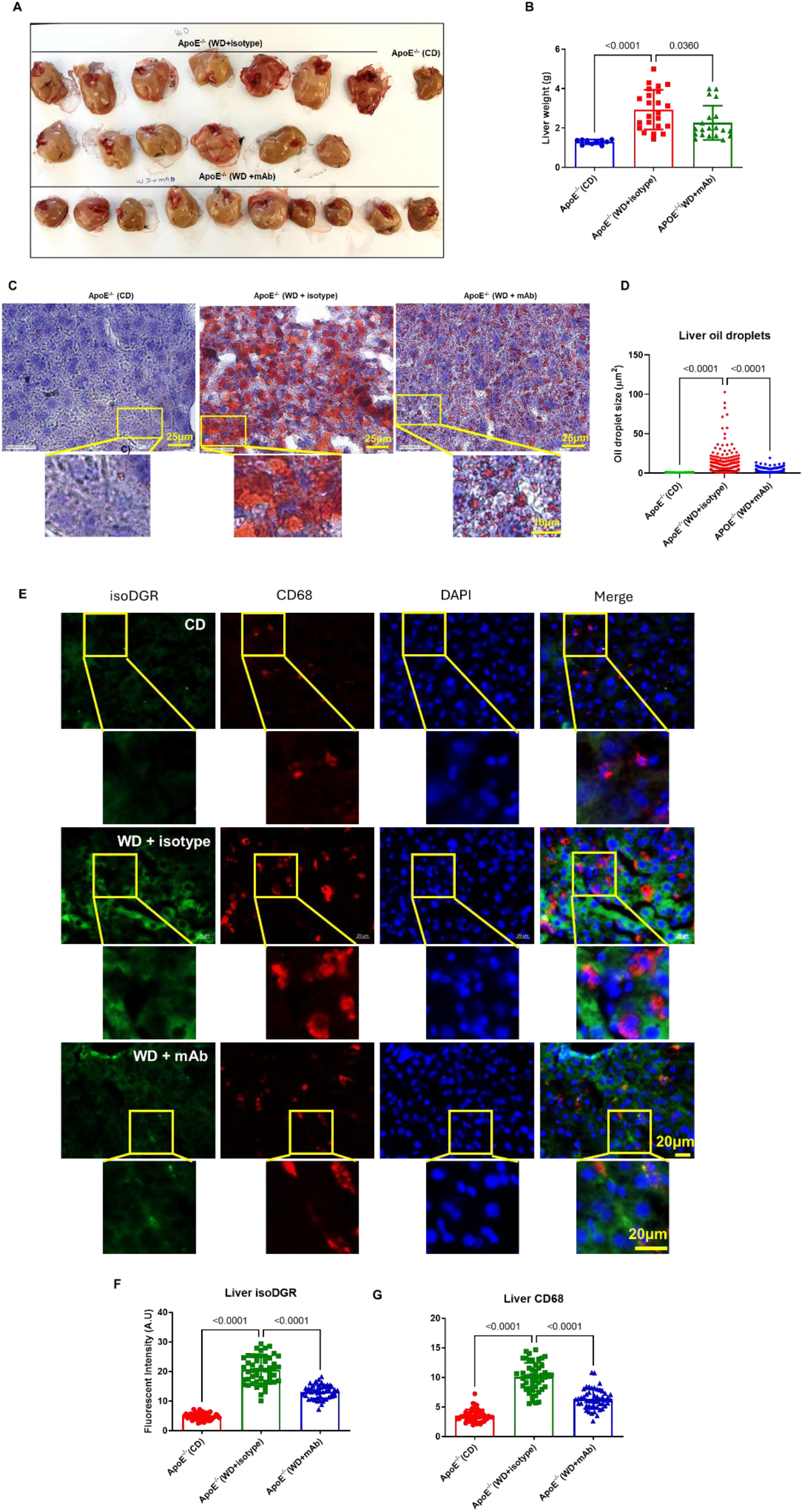
IsoDGR-mAb attenuates diet-induced fatty liver disease and hepatic inflammation. (A) Gross liver images from ApoE^-/-^ mice after 16 weeks: CD-fed (right), WD-fed receiving isotype control (first two rows), and WD-fed treated with isoDGR-mAb (third row). WD causes hepatomegaly and pale, fatty livers, whereas isoDGR-mAb treatment results in smaller livers with healthier appearance. (B) Liver weight for the same groups. WD feeding significantly increased liver mass (steatosis), while isoDGR-mAb treatment significantly reduced liver weight. (C) Representative liver sections stained with Oil Red O. Chow diet liver shows minimal lipid accumulation (almost no red staining), WD control liver is filled with lipid droplets (extensive red), and WD+mAb liver has markedly lower lipid content. (D) Quantification of hepatic lipid area (% Oil Red O+ area) shows that WD+mAb livers contain >60% less lipid than WD controls. (E) Liver immunofluorescence for isoDGR (green) and macrophages (CD68, red). WD control liver exhibits extensive isoDGR deposition and numerous CD68+ Kupffer cells/macrophages, whereas the WD+mAb liver shows greatly diminished isoDGR and fewer CD68+ cells. Quantification of (F) isoDGR fluorescence intensity and (G) CD68+ cell area in livers confirms significant reductions of both upon isoDGR-mAb treatment. Thus, targeting isoDGR protects against WD-induced hepatic steatosis and inflammation. Each experimental group included 6-12 mice (n = 6-12).

We then assessed isoDGR deposition in the liver, which was strongly associated with local inflammation (Figure 5E): WD-fed control mice displayed significant isoDGR-positive staining along hepatic sinusoids as well as numerous CD68+ liver macrophages (Kupffer cells and infiltrating monocytes). IsoDGR-mAb treatment greatly reduced isoDGR staining in the liver and concomitantly decreased the frequency of CD68+ macrophages. Quantitative analysis confirmed that hepatic isoDGR signal (Figure 5F) and macrophage infiltration (Figure 5G) were lower in treated mice than in untreated WD mice (*p* < 0.0001 for both). These data suggest that in WD-fed *ApoE*^-/-^ mice, isoDGR-modified proteins accumulate in the liver and contribute to inflammatory responses via recruitment of monocyte-macrophages. Conversely, antibody neutralisation of isoDGR epitopes inhibits monocyte recruitment and activation of Kupffer cells, leading to decreased hepatic inflammation. The combined effect of these changes is that isoDGR-mAb-treated mice were largely protected from WD-induced steatohepatitis.

### IsoDGR neutralization boosts metabolic rate

To understand how isoDGR-mAb treatment reduces fat accumulation in liver tissue, we conducted metabolic cage experiments to assess mouse metabolism after 16 weeks WD feeding and 8 weeks antibody treatment. Despite similar food intake and physical activity, WD-fed mice treated with isoDGR-mAb displayed increased energy expenditure relative to controls, with indirect calorimetry indicating ∼30% higher oxygen consumption (VO_2_) and carbon dioxide production across both light and dark cycles (Figure 6A, B). These data suggest an elevated basal metabolic rate in isoDGR-mAb-treated mice, which were burning significantly more calories than untreated animals despite identical diets in both groups.

**Figure 6.**
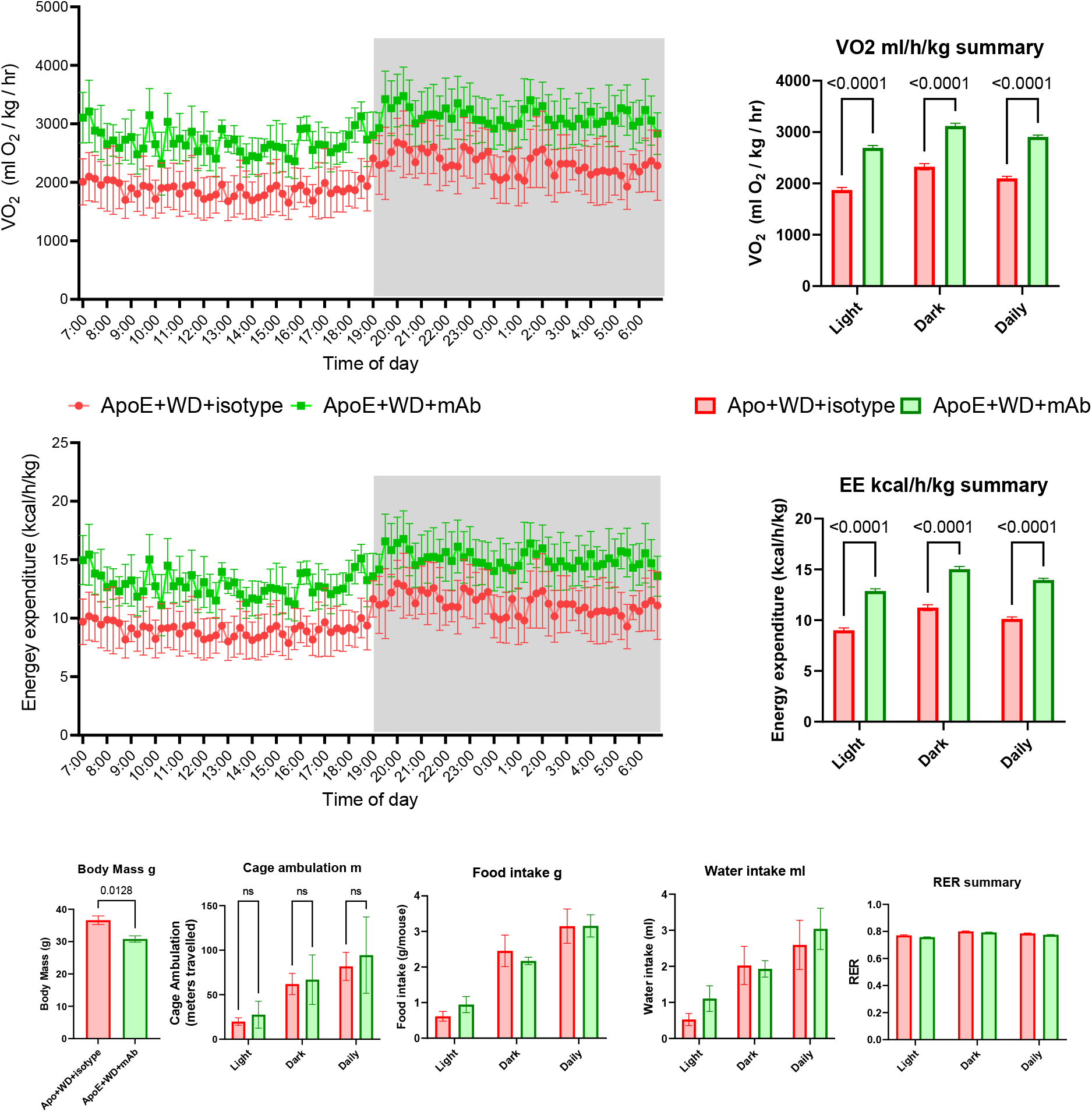
IsoDGR-mAb increases energy expenditure without change in food intake or activity. Metabolic cage analysis of WD-fed ApoE^-/-^ mice with or without isoDGR-mAb treatment. (A) Hourly oxygen consumption (VO_2_) over 48 hours (dark phase shaded) for isotype-treated (red) vs isoDGR-mAb-treated mice (green). Treated mice displayed consistently higher VO_2_throughout. (B) Average VO_2_in light and dark phases (as well as 24 hours total) was ∼30% higher in isoDGR-mAb-treated mice vs controls (p < 0.0001). (C) Carbon dioxide production and calculated energy expenditure (EE) also ran higher in treated mice across the day. (D) Mean EE in light phase, dark phase, and over 24 hours confirmed a significant increase of ∼30% with isoDGR-mAb treatment. (E) Final body weights of mice after 16 weeks: isoDGR-mAb-treated mice weighed significantly less than controls (p < 0.05). (F) Total locomotor activity (distance traveled) in light and dark periods is similar between treated and control mice. (G, H) Daily food intake and water intake show no significant differences between groups. (I) Respiratory exchange ratio (RER = VCO_2_/VO_2_) remained at ∼0.75 for both groups, indicating comparable substrate utilization. Together, these data indicate that isoDGR neutralization enhances basal metabolic rate and prevents excess weight gain, likely by relieving inflammation-induced metabolic suppression (rather than by significantly affecting appetite or activity). Each experimental group included 4 mice (n = 4).

Importantly, the increased metabolism observed following isoDGR-mAb treatment was not due to differences in behavior or caloric intake. Locomotor activity was monitored and showed only a marginal increase in the isoDGR-mAb-treated group (p = ns; Figure 6F). Both WD isotype- and WD isoDGR-treated mice exhibited similar activity patterns and total movement counts during both light (low activity) and dark (high activity) phases. This finding indicates that treated mice were not simply more active, but instead expended more energy at a given activity level. Similarly, food and water intake over 48 hours were nearly identical between the two groups (Figure 6G, H). Treated mice and control mice consumed comparable amounts of calorie-dense WD, indicating that the increased energy expenditure (EE) could not be attributed to changes in caloric intake. Additionally, the respiratory exchange ratio (RER) remained at ∼0.75 in both groups (Figure 6I), consistent with predominant fat oxidation, as expected for mice on high-fat diet. The unchanged RER suggests that fuel substrate preference (fat vs. carbohydrate utilization) was similar in both groups. Thus, isoDGR-mAb did not cause a shift in fuel type, but rather increased the total amount of fuel oxidized over a given period. Together, these findings suggest that increased energy expenditure with isoDGR-mAb treatment is not due to significant behavioral changes and more likely reflects a direct metabolic impact of antibody treatment, potentially due to reduced inflammatory burden.

## Discussion

Our findings establish the isoDGR motif as a critical mediator of atherosclerosis and metabolic dysfunction in the context of high-fat diet. In both an accelerated protein damage model (*Pcmt1*^-/-^ mice) and a diet-induced disease model (*ApoE*^-/-^ mice fed WD), isoDGR accumulation correlated with vascular tissue damage and inflammation, whereas antibody neutralization of isoDGR induced a range of marked physiological improvements. These findings indicate a potentially new therapeutic approach by targeting specific molecular damage motifs that accumulate with aging. By neutralising an age-related neo-epitope such as isoDGR, it is possible to remove a key catalyst of tissue degeneration and chronic inflammation, thereby mitigating downstream pathology.

Mechanistically, our results align with the concept that isoDGR-modified matrix proteins in the vasculature can act as pro-inflammatory ligands. These neo-epitopes can bind integrins on monocytes, macrophages, and endothelial cells, leading to activation of pro-inflammatory pathways and promoting leukocyte recruitment to the vessel wall. In the untreated *ApoE*^-/-^ mice, we observed strong correlation of isoDGR with infiltrating macrophages in plaque tissue, similar to previous observations that isoDGR-modified fibronectin correlates with CD68+ macrophage frequency in human plaques ^3, 4^. Such interactions likely create a vicious cycle: integrin engagement by isoDGR triggers macrophages to secrete cytokines and proteases, leading to further matrix remodeling and exposure of additional damage motifs. Results in the current report indicate that anti-isoDGR therapy can break this cycle, since treated mice had fewer lesional macrophages and lower concentrations of systemic CRP and inflammatory cytokines.

Beyond the vascular wall effects, an intriguing outcome of isoDGR targeting was the enhancement of metabolic performance. Treated mice displayed increased oxygen consumption and resistance to fatty liver disease despite unaltered caloric intake. This phenomenon is consistent with emerging evidence that chronic inflammation acts as a brake on energy expenditure, whereas decreasing inflammatory stress can reactivate thermogenic and metabolic pathways ^22-24^. By dampening inflammation via removal of isoDGR damage cues, the antibody likely enabled higher basal metabolism, as reflected by elevated VO_2_ and reduced steatosis. This result indicates that therapies addressing molecular damage can confer systemic benefits beyond the primary disease (atherosclerosis) and improve overall metabolic health.

Collectively, our study provides an important proof-of-concept that immunotherapeutic targeting of age-linked protein damage motifs can simultaneously combat cardiovascular and metabolic components of disease. These findings extend the emerging concept that ‘damage-clearing’ therapies can restore tissue homeostasis by removing pathological protein modifications and senescent cells ^9, 18, 25-29^. Notably, isoDGR is just one class of many spontaneous damage proteins that accumulate with aging, hence other classes of DPMs could potentially also be targeted in similar fashion (e.g. carbamylation, thiol trioxidation, carboxyethyl-lysine, or other advanced glycation/oxidation products)^30-39^. Our results strongly suggest that the scope of immunotherapeutic targets can now be extended to include endogenous degeneration-associated antigens, not just cytokines, pathogens, and tumor biomolecules.

Looking forward, our observations also suggest that isoDGR or related damage motifs serve as biomarkers for cardiovascular risk stratification in aging populations^2, 4^. Therapies such as isoDGR-mAb could perhaps be developed to complement existing treatments (e.g., statins or anti-inflammatory drugs) by specifically addressing age-linked elements of disease. Further preclinical studies are now warranted to evaluate long-term safety, optimal timing of treatment (e.g. mid-life intervention to prevent isoDGR accumulation), and effects in other models of cardiovascular aging and injury. Ultimately, clinical trials will be needed to determine if targeting isoDGR can slow atherosclerosis progression and improve health-span in humans. Our findings lay the groundwork for a novel class of ‘anti-degenerative’ therapies that target the molecular byproducts of aging to treat atherosclerosis underlying cardiovascular disease.

## References

1. Sen I, Trzaskalski NA, Hsiao YT, Liu PP, Shimizu I and Derumeaux GA. Aging at the Crossroads of Organ Interactions: Implications for the Heart. Circ Res. 2025:1286–1305.

2. Mol BM, Motsak T, Hoekstra JKR, Zuluaga VO, Rumpff-Derksen S, Paspali-Strik I, Ngan SC, Pasterkamp G, de Borst GJ, Sze SK and de Kleijn DPV. High Levels of Plasma and Plaque IsoDGR-Modified Fibronectin are Associated with Rupture Prone Plaque Characteristics in Carotid Endarterectomy Patients. Eur J Vasc Endovasc Surg.

3. Dutta B, Park JE, Kumar S, Hao P, Gallart-Palau X, Serra A, Ren Y, Sorokin V, Lee CN, Ho HH, de Kleijn D and Sze SK. Monocyte adhesion to atherosclerotic matrix proteins is enhanced by Asn-Gly-Arg deamidation. Sci Rep. 2017:5765.

4. Park JE, JebaMercy G, Pazhanchamy K, Guo X, Ngan SC, Liou KCK, Lynn SE, Ng SS, Meng W, Lim SC, Leow MK, Richards AM, Pennington DJ, de Kleijn DPV, Sorokin V, Ho HH, McCarthy NE and Sze SK. Aging-induced isoDGR-modified fibronectin activates monocytic and endothelial cells to promote atherosclerosis. Atherosclerosis. 2021:58–68.

5. Cheow ESH, Hao PL, Hao PL, Sorokin V, Lee CN, Kleijn DPV and Sze SK. The Role of Protein Deamidation in Cardiovascular Disease. Proceedings of the 23rd American Peptide Symposium. 2013;17:212–213.

6. Spitaleri A, Mari S, Curnis F, Traversari C, Longhi R, Bordignon C, Corti A, Rizzardi GP and Musco G. Structural basis for the interaction of isoDGR with the RGD-binding site of alphavbeta3 integrin. J Biol Chem. 2008:19757–68.

7. Curnis F, Longhi R, Crippa L, Cattaneo A, Dondossola E, Bachi A and Corti A. Spontaneous formation of L-isoaspartate and gain of function in fibronectin. J Biol Chem. 2006:36466–76.

8. Park JE, JebaMercy G, Pazhanchamy K, Guo X, Ngan SC, Liou KCK, Lynn SE, Ng SS, Meng W, Lim SC, Leow MK, Richards AM, Pennington DJ, de Kleijn DPV, Sorokin V, Ho HH, McCarthy NE and Sze SK. Aging-induced isoDGR-modified fibronectin activates monocytic and endothelial cells to promote atherosclerosis. Atherosclerosis. 2021;324:58–68.

9. Kalailingam P, Mohd-Kahliab KH, Ngan SC, Iyappan R, Melekh E, Lu T, Zien GW, Sharma B, Guo T, MacNeil AJ, MacPherson RE, Tsiani EL, O’Leary DD, Lim KL, Su IH, Gao YG, Richards AM, Kalaria RN, Chen CP, McCarthy NE and Sze SK. Immunotherapy targeting isoDGR-protein damage extends lifespan in a mouse model of protein deamidation. EMBO molecular medicine. 2023;15:e18526.

10. Kalailingam P, Ngan SC, Iyappan R, Nehchiri A, Mohd-Kahliab KH, Lee BST, Sharma B, Machan R, Bo ST, Chambers ES, Fajardo VA, Macpherson REK, Liu J, Klentrou P, Tsiani EL, Lim KL, Su IH, Gao YG, Richar AM, Kalaria RN, Chen CP, Balion C, de Kleijn D, McCarthy NE and Sze SK. Immunotherapeutic targeting of aging-associated isoDGR motif in chronic lung inflammation. Aging Cell. 2025:e14425.

11. Curnis F, Sacchi A, Gasparri A, Longhi R, Bachi A, Doglioni C, Bordignon C, Traversari C, Rizzardi GP and Corti A. Isoaspartate-glycine-arginine: a new tumor vasculature-targeting motif. Cancer Res. 2008:7073–82.

12. Dutta B, Park JE, Kumar S, Hao P, Gallart-Palau X, Serra A, Ren Y, Sorokin V, Lee CN, Ho HH, de Kleijn D and Sze SK. Monocyte adhesion to atherosclerotic matrix proteins is enhanced by Asn-Gly-Arg deamidation. Scientific reports. 2017;7:5765.

13. Wang J, Xia J, Su J, Cao Z, Yang W, Zhang P and Xu Y. Multi-omics Analysis Sheds Light on the Extracellular Role of PCMT1. J Proteome Res.

14. Xia J, Hou Y, Wang J, Zhang J, Wu J, Yu X, Cai H, Yang W, Xu Y and Mou S. Repair of Isoaspartyl Residues by PCMT1 and Kidney Fibrosis. J Am Soc Nephrol.

15. Kim E, Lowenson JD, Clarke S and Young SG. Phenotypic analysis of seizure-prone mice lacking L-isoaspartate (D-aspartate) O-methyltransferase. J Biol Chem. 1999:20671–8.

16. Wang H, Chang TS, Dombroski BA, Cheng PL, Patil V, Valiente-Banuet L, Farrell K, McLean C, Molina-Porcel L, Rajput A, De Deyn PP, Le Bastard N, Gearing M, Kaat LD, Van Swieten JC, Dopper E, Ghetti BF, Newell KL, Troakes C, de Yébenes JG, Rábano-Gutierrez A, Meller T, Oertel WH, Respondek G, Stamelou M, Arzberger T, Roeber S, Müller U, Hopfner F, Pastor P, Brice A, Durr A, Le Ber I, Beach TG, Serrano GE, Hazrati LN, Litvan I, Rademakers R, Ross OA, Galasko D, Boxer AL, Miller BL, Seeley WW, Van Deerlin VM, Lee EB, White CL, 3rd, Morris H, de Silva R, Crary JF, Goate AM, Friedman JS, Leung YY, Coppola G, Naj AC, Wang LS, Dalgard C, Dickson DW, Höglinger GU, Schellenberg GD, Geschwind DH and Lee WP. Whole-genome sequencing analysis reveals new susceptibility loci and structural variants associated with progressive supranuclear palsy. Mol Neurodegener. 2024:61.

17. Kim E, Lowenson JD, MacLaren DC, Clarke S and Young SG. Deficiency of a protein-repair enzyme results in the accumulation of altered proteins, retardation of growth, and fatal seizures in mice. Proceedings of the National Academy of Sciences of the United States of America. 1997;94:6132–7.

18. Kalailingam P, Ngan SC, Iyappan R, Nehchiri A, Mohd-Kahliab KH, Lee BST, Sharma B, Machan R, Bo ST, Chambers ES, Fajardo VA, Macpherson REK, Liu J, Klentrou P, Tsiani EL, Lim KL, Su IH, Gao YG, Richar AM, Kalaria RN, Chen CP, Balion C, de Kleijn D, McCarthy NE and Sze SK. Immunotherapeutic targeting of aging-associated isoDGR motif in chronic lung inflammation. Aging Cell. 2025;24:e14425.

19. Kwiatkowski A, Co C, Kameoka S, Zhang A, Coughlin J, Cameron T, Chiao E, Bergelson S and Schmid Mason C. Assessment of the role of afucosylated glycoforms on the in vitro antibody-dependent phagocytosis activity of an antibody to Aβ aggregates. MAbs. 2020;12:1803645.

20. Ilyas I, Little PJ, Liu Z, Xu Y, Kamato D, Berk BC, Weng J and Xu S. Mouse models of atherosclerosis in translational research. Trends Pharmacol Sci. 2022:920–939.

21. Zhang SH, Reddick RL, Piedrahita JA and Maeda N. Spontaneous hypercholesterolemia and arterial lesions in mice lacking apolipoprotein E. Science. 1992;258:468–71.

22. Yan S, Kumari M, Xiao H, Jacobs C, Kochumon S, Jedrychowski M, Chouchani E, Ahmad R and Rosen ED. IRF3 reduces adipose thermogenesis via ISG15-mediated reprogramming of glycolysis. J Clin Invest. 2021;131.

23. Zhu Y, Yang R, Deng Z, Deng B, Zhao K, Dai C, Wei G, Wang Y, Zheng J, Ren Z, Lv W, Xiao Y, Mei Z and Song T. Adipose Tissue-Resident Sphingomonas Paucimobilis Suppresses Adaptive Thermogenesis by Reducing 15-HETE Production and Inhibiting AMPK Pathway. Adv Sci (Weinh). 2024:e2310236.

24. Wu D, Eeda V, Maria Z, Rawal K, Wang A, Herlea-Pana O, Babu Undi R, Lim HY and Wang W. Targeting IRE1α improves insulin sensitivity and thermogenesis and suppresses metabolically active adipose tissue macrophages in male obese mice. Elife. 2025.

25. Aguado J, Amarilla AA, Taherian Fard A, Albornoz EA, Tyshkovskiy A, Schwabenland M, Chaggar HK, Modhiran N, Gómez-Inclán C, Javed I, Baradar AA, Liang B, Peng L, Dharmaratne M, Pietrogrande G, Padmanabhan P, Freney ME, Parry R, Sng JDJ, Isaacs A, Khromykh AA, Valenzuela Nieto G, Rojas-Fernandez A, Davis TP, Prinz M, Bengsch B, Gladyshev VN, Woodruff TM, Mar JC, Watterson D and Wolvetang EJ. Senolytic therapy alleviates physiological human brain aging and COVID-19 neuropathology. Nat Aging. 2023:1561–1575.

26. van Olst L, Simonton B, Edwards AJ, Forsyth AV, Boles J, Jamshidi P, Watson T, Shepard N, Krainc T, Argue BM, Zhang Z, Kuruvilla J, Camp L, Li M, Xu H, Norman JL, Cahan J, Vassar R, Chen J, Castellani RJ, Nicoll JA, Boche D and Gate D. Microglial mechanisms drive amyloid-β clearance in immunized patients with Alzheimer’s disease. Nat Med. 2025;31:1604–1616.

27. Jeon OH, Kim C, Laberge RM, Demaria M, Rathod S, Vasserot AP, Chung JW, Kim DH, Poon Y, David N, Baker DJ, van Deursen JM, Campisi J and Elisseeff JH. Local clearance of senescent cells attenuates the development of post-traumatic osteoarthritis and creates a pro-regenerative environment. Nat Med. 2017;23:775–781.

28. Chang J, Wang Y, Shao L, Laberge RM, Demaria M, Campisi J, Janakiraman K, Sharpless NE, Ding S, Feng W, Luo Y, Wang X, Aykin-Burns N, Krager K, Ponnappan U, Hauer-Jensen M, Meng A and Zhou D. Clearance of senescent cells by ABT263 rejuvenates aged hematopoietic stem cells in mice. Nat Med. 2016;22:78–83.

29. Pluvinage JV, Haney MS, Smith BAH, Sun J, Iram T, Bonanno L, Li L, Lee DP, Morgens DW, Yang AC, Shuken SR, Gate D, Scott M, Khatri P, Luo J, Bertozzi CR, Bassik MC and Wyss-Coray T. CD22 blockade restores homeostatic microglial phagocytosis in ageing brains. Nature. 2019;568:187–192.

30. Gallart-Palau X, Tan LM, Serra A, Gao Y, Ho HH, Richards AM, Kandiah N, Chen CP, Kalaria RN and Sze SK. Degenerative protein modifications in the aging vasculature and central nervous system: A problem shared is not always halved. Ageing Res Rev. 2019:100909.

31. Sánchez Milán JA, Fernández-Rhodes M, Guo X, Mulet M, Ngan SC, Iyappan R, Katoueezadeh M, Sze SK, Serra A and Gallart-Palau X. Trioxidized cysteine in the aging proteome mimics the structural dynamics and interactome of phosphorylated serine. Aging Cell. 2024;23:e14062.

32. Adav SS and Sze SK. Hypoxia-Induced Degenerative Protein Modifications Associated with Aging and Age-Associated Disorders. Aging Dis. 2020;11:341–364.

33. Gallart-Palau X, Serra A, Lee BST, Guo X and Sze SK. Brain ureido degenerative protein modifications are associated with neuroinflammation and proteinopathy in Alzheimer’s disease with cerebrovascular disease. J Neuroinflammation. 2017:175.

34. Paramasivan S, Adav SS, Ngan SC, Dalan R, Leow MK, Ho HH and Sze SK. Serum albumin cysteine trioxidation is a potential oxidative stress biomarker of type 2 diabetes mellitus. Sci Rep. 2020:6475.

35. Adav SS and Sze SK. Insight of brain degenerative protein modifications in the pathology of neurodegeneration and dementia by proteomic profiling. Mol Brain. 2016:92.

36. Gallart-Palau X, Serra A and Sze SK. Uncovering Neurodegenerative Protein Modifications via Proteomic Profiling. Int Rev Neurobiol. 2015:87–116.

37. Gladyshev VN, Kritchevsky SB, Clarke SG, Cuervo AM, Fiehn O, de Magalhães JP, Mau T, Maes M, Moritz R, Niedernhofer LJ, Van Schaftingen E, Tranah GJ, Walsh K, Yura Y, Zhang B and Cummings SR. Molecular Damage in Aging. Nat Aging. 2021:1096–1106.

38. Zhang Y, Unnikrishnan A, Deepa SS, Liu Y, Li Y, Ikeno Y, Sosnowska D, Van Remmen H and Richardson A. A new role for oxidative stress in aging: The accelerated aging phenotype in Sod1(−/)(−) mice is correlated to increased cellular senescence. Redox Biol. 2017:30–37.

39. Baldensperger T, Preissler M and Becker CFW. Non-enzymatic posttranslational protein modifications in protein aggregation and neurodegenerative diseases. RSC Chem Biol. 2025;6:129–149.

